# Individual prediction tendencies facilitate cortical speech tracking

**DOI:** 10.1101/2022.04.22.489224

**Authors:** Juliane Schubert, Fabian Schmidt, Quirin Gehmacher, Annika Bresgen, Nathan Weisz

## Abstract

Listening can be conceptualized as a process of active inference, in which the brain forms internal models to predict and integrate auditory information in a complex interaction of bottom-up and top-down processes. Whether inter-individual “prediction tendencies” shape listening experiences of real-world stimuli such as speech is, however, unknown. In the current study, we used a passive paradigm presenting tone sequences of varying entropy level, to independently quantify auditory prediction tendency (as the tendency to anticipate low-level acoustic features according to their contextual probability) for each individual. This measure was then used to predict the magnitude of cortical speech (envelope) tracking in a multi speaker listening task, where participants listened to audiobooks narrated by a target speaker in isolation or interfered by 1 or 2 distractors. Furthermore, rare semantic violations were introduced into the story, enabling us to also examine effects of word surprisal during continuous speech processing. Our results show that individual prediction tendency facilitates cortical speech tracking. Furthermore, we find interactions between individual prediction tendency and background noise as well as word surprisal in disparate brain regions. In sum, our findings suggest that individual prediction tendencies are generalizable across different listening situations and may serve as a valuable element to explain interindividual differences in natural listening experience.

## INTRODUCTION

Listening is a neurobiological challenge that requires a complex interaction of bottom-up and top-down processes. For instance, understanding speech in a noisy environment by bottom-up input alone would be impossible due to the vast amount of overlapping spectrotemporal information (McDermott, 2009). In line with notions of the so-called predictive brain (K. Friston, 2010; Knill & Pouget, 2004; Yon et al., 2019), we assume that our brain is actively engaged when listening to speech by fitting and testing internal models, inferring which sound sources (“auditory objects”; Griffiths & Warren, 2004) are causing the neural activity patterns. This process requires the constant generation of predictions that are continuously compared with incoming bottom-up information.

With the rising scientific interest to prove the ubiquity of the predictive brain, also the consideration of individual differences has received increased attention. In search of a predictor of linguistic abilities, individual differences in statistical learning, “a general capacity for picking up regularities”, have been proposed in this regard (for a review see Siegelman et al., 2017). Crucially, however, statistical learning as an individual capacity should be operationalised and precisely which of its components drive the relationship between statistical learning and linguistic performance remains unclear. We propose that the individual tendency to actively generate auditory predictions might be one such factor. Following up on research focusing on individual differences, it is reasonable to assume that humans differ in their capability as well as their overall tendency to actively predict incoming sensory input. These differences have already been linked to a variety of clinical psychological conditions and disorders such as autism (Sinha et al., 2014), schizophrenia (Sterzer et al., 2018) and tinnitus (Partyka et al., 2019; Sedley et al., 2016). However, prediction tendencies have so far not been studied as a potential factor that drives interindividual differences in everyday listening abilities such as speech. Friston and colleagues (2021) recently provide a theoretical perspective in which the predictive brain is engaged in active listening, for example supporting (covertly) the dissection of words from a continuous auditory signal based on prior assumptions. Here we aim to consider the influence of individual differences in prediction tendencies on speech processing. Our main assumption is that individual prediction tendency, which we operationalize as the tendency to pre-activate sensory features of strong likelihood, contributes substantially to differences in speech processing.

Another goal of the current study is to detail the circumstances under which strong prediction tendencies are particularly beneficial for speech processing. Indeed, “normal” hearing individuals vary considerably in their ability to understand speech with some even reporting difficulties in everyday speech comprehension and communication (Ruggles et al., 2011). These interindividual differences become especially apparent in noisy situations which are typical for natural environments. In the iconic cocktail party situation multiple speech streams enter the auditory system, yet only one is to be followed (Oberfeld & Klöckner-Nowotny, 2016). This task gets increasingly challenging with increased interfering noise, revealing and/or increasing inter-individual differences in listening experiences (Wöstmann et al., 2015). To establish these contextual influences on speech processing with high temporal resolution, cortical tracking of the acoustic envelope (extracted from the audiofile and used as a predictor to explain variance in the neural signal) has become increasingly popular (Brodbeck et al., 2018; Vanthornhout et al., 2018).

Regardless of signal-to-noise ratio, it should be considered that even under optimal conditions not all incoming sensory information is equally predictable. Thus, how the influence of prediction tendency on speech processing depends on the predictability of the speech input itself is another important question that needs to be addressed. Although there are several neuroimaging studies offering valuable insight to the effect of prior knowledge on isolated sentences (Di Liberto et al., 2018) or target words (Sohoglu et al., 2012), it has been questioned whether effects from controlled manipulation generalize to natural, narrative language. Using artificial neural network (ANN) language models (which mimic human language processing; Schrimpf et al., 2021) to quantify word predictability on a continuous scale, recent evidence suggests cortical tracking of surprisal during continuous speech perception (Donhauser & Baillet, 2020; Weissbart et al., 2020). Broderick and colleagues (2019) further showed that semantic predictability also enhances acoustic envelope encoding in continuous speech. Although the benefits of using natural speech with no experimentally induced manipulations have to be acknowledged, one remaining disadvantage is the difficulty in separating top-down from bottom-up influences. In narrative language the acoustic signal is correlated with the higher level information it conveys and any alteration in speech envelope encoding may be attributed to both aspects (Brodbeck & Simon, 2020).

In the current study, we combined magnetoencephalographic (MEG) data from two different paradigms in order to establish a link between individual prediction tendency and speech perception: 1) An adapted paradigm as used by Demarchi and colleagues (2019), presenting tone sequences of varying entropy level (in the following referred to as entropy modulation paradigm) which was used to independently quantify individual prediction tendency. 2) A multi speaker listening task to investigate individual differences in neural speech tracking across different levels of background noise. We used natural, continuous speech but also introduced rare semantic violations by randomly replacing a limited set of words with target words that occurred elsewhere in the same storyline. This allowed us to investigate encoding of the acoustic envelope on a continuous scale as well as on an individual word level (of lexically identical stimuli), ensuring that effects of word surprisal cannot be attributed to differences in bottom-up input. Assuming that a strong individual prediction tendency will allow for higher experiential certainty in language segments with low surprise, we expected strong contextual violations to interfere with this facilitation of speech processing. Our results show that prediction tendency, which we define as the tendency to anticipate auditory events of high probability, facilitates cortical speech envelope encoding in perisylvian and inferior frontal areas of both hemispheres. Furthermore, we find spatially dissociable interactions between individual prediction tendency and background noise as well as word surprisal. In sum, these findings suggest that individual prediction tendencies show considerable generalizability across different listening situations and may serve as a valuable element to explain interindividual differences in natural listening experience.

## METHODS

### Subjects

In total 53 (21 male) subjects were recruited to participate in the study, however 4 subjects did not complete the experiment due to technical difficulties, leaving a total number of 49 subjects (mean age = 26.31, range = 19 - 40) for further data analysis. Participation was compensated either monetarily or via course credits. All participants were German native speakers and reported normal hearing which was confirmed using standard pure tone audiometry. Participants gave written, informed consent and reported that they had no previous neurological or psychiatric disorders. The experimental procedure was approved by the ethics committee of the University of Salzburg and was carried out in accordance with the declaration of Helsinki.

### Stimuli

We used audio recordings of excerpts from German books and short-stories (see Appendix table 1). In total, 6 different, consistent target stories (à ∼10min) and 18 distractor stories (à ∼3min) were recorded with a t.bone SC 400 studio microphone and a sampling rate of 44100 Hz. Prior to recording, target stories were split into separate trials of approximately 3 - 4 min (mean = 3.46 min, range = 3.05 - 4.07 min). In addition, we randomly selected half of the nouns that ended a sentence and replaced them with the other half to induce unexpected semantic violations within each trial, resulting in two sets of lexically identical words (N = 79) that differed greatly in their contextual probabilities (see **Fig. 2C** for an example). All 6 target stories were recorded twice, narrated by a different speaker (male vs. female). The remaining 18 recordings were narrated by the same two speakers (used as second female/male distractor speaker to a male/female target speaker respectively) and two additional speakers (used as first distractor speaker). Stimuli were presented in 6 blocks containing 3 trials each, resulting in 3 male and 3 female target speaker blocks for every participant (see next section).

### Experimental procedure

Before the start of the experiment, we performed standard pure tone audiometry using the AS608 Basic (Interacoustics, Middelfart, Denmark) in order to assess participants’ individual hearing ability. Afterwards, participants’ individual head shapes were assessed using cardinal head points (nasion and pre-auricular points), digitized with a Polhemus Fastrak Digitizer (Polhemus) and around 300 points on the scalp. For every participant MEG sessions started with a 5-minute resting state recording, after which the individual hearing threshold was determined using a pure tone of 1043 Hz. This was followed by 2 blocks of passive listening to tone sequences of varying entropy level to quantify individual prediction tendencies (see quantification of individual prediction tendency) while participants watched a landscape movie (LoungeV Films, 2017). In the main task 6 different stories were presented in separate blocks in random order and with randomly balanced selection of the target speaker (male vs. female voice). Each block consisted of 3 trials with a continuous storyline, with each trial corresponding to one of 3 experimental conditions: a single speaker only, a 1-distractor speaker and a 2-distractor speaker condition (in the following: 0-dist, 1-dist, 2-dist, see **Fig. 2A**). The distractor speakers were always selected to be of the opposite sex of the target speaker and were presented exactly 20 s after target speaker onset. All stimuli were presented binaurally at equal volume for the left and right ear (i.e. at phantom center). Participants were instructed to always attend to the first speaker and their understanding was tested using comprehension questions (true vs. false statements) at the end of every trial. Further, participants indicated their intrinsic motivation and their perceived task-difficulty on a 5-point likert scale at the end of every trial. All stimuli were presented at 40db above the individual hearing threshold. In total, the experiment lasted approximately 3.5h per participant (including MEG preparation time). The experiment was coded and conducted with the Psychtoolbox-3 (Brainard, 1997; Kleiner et al., 2007) with an additional class-based library (‘Objective Psychophysics Toolbox’, o_ptb) on top of it (Hartmann & Weisz, 2020).

### MEG Data Acquisition and Analysis

A whole head MEG system (Elekta Neuromag Triux, Elekta Oy, Finland), placed within a standard passive magnetically shielded room (AK3b, Vacuumschmelze, Germany), was used to capture magnetic brain activity with a sampling frequency of 10 kHz (hardware filters: 0.1 - 3300 Hz). The signal was recorded with 102 magnetometers and 204 orthogonally placed planar gradiometers at 102 different positions. In a first step, a signal space separation algorithm, implemented in the Maxfilter program (version 2.2.15) provided by the MEG manufacturer, was used to clean the data from external noise and realign data from different blocks to a common standard head position. Data preprocessing was performed using Matlab R2020b (The MathWorks, Natick, Massachusetts, USA) and the FieldTrip Toolbox (Oostenveld et al., 2010). All data was filtered between 0.1 Hz and 30 Hz (Kaiser windowed finite impulse response filter) and downsampled to 100 Hz. To identify eye-blinks and heart rate artifacts, 50 independent components were identified from filtered (0.1 - 100 Hz), downsampled (1000 Hz) continuous data of the recordings from the entropy modulation paradigm and on average 3 components were removed for every subject. Data of the entropy modulation paradigm was epoched into segments of 1200 ms (from 400 ms before sound onset to 800 ms after onset). Multivariate pattern analysis (see quantification of individual prediction tendency) was carried out using the MVPA-Light package (Treder, 2020). Single trial data from the main task were further projected into source-space using LCMV spatial filters (Van Veen et al., 1997). To compute these filters, anatomical template images were warped to the individual head shape and brought into a common space by co-registering them based on the three anatomical landmarks (nasion, left and right preauricular points) with a standard brain from the Montreal Neurological Institute (MNI, Montreal, Canada; (Mattout et al., 2007)). Afterwards, a single-shell head model (Nolte, 2003) was computed for each participant. As a source model, a grid with 1 cm resolution and 2982 voxels based on an MNI template brain was morphed into the brain volume of each participant. This allows group-level averaging and statistical analysis as all the grid points in the warped grid belong to the same brain region across subjects. Afterwards, the data was temporally aligned with the corresponding features from the audio material.

### Quantification of individual prediction tendency

In order to quantify individual prediction tendency, we used an entropy modulation paradigm (see also Demarchi et al., 2019) where participants passively listened to sequences of 4 different pure tones (1: 440 Hz, 2: 587 Hz, 3: 782 Hz, 4: 1043 Hz, each lasting 100 ms) during two separate blocks, each consisting of 1500 tones presented with a temporally predictable rate of 3 Hz. Transitional probabilities between tones varied across two different levels of entropy (ordered vs. random; see **Fig. 1A**). While in an “ordered” context certain transitions (hereinafter referred to as forward transitions, i.e. 1→2, 2→3, 3→4, 4→1) were to be expected with a high probability of 75%, self repetitions (e.g., 1→1, 2→2,…) were rather unlikely with a probability of 25%. However, in a “random” context all possible transitions were equally likely with a probability of 25%. Entropy levels changed between the two blocks for half of the participants and pseudorandomly every 500 trials within each block for the other half, always resulting in a total of 1500 trials per entropy condition. As Demarchi et al. (2019) showed that preactivation of carrier-frequency related neural activity is systematically enhanced within low entropy conditions, it can be assumed that correct frequency-specific predictions about upcoming events should only be generated for transitions of high probability, i.e. (expected) forward transitions in an ordered context. Whereas in the same ordered context, unexpected self-repetition trials should erroneously generate prestimulus predictions to an upcoming tone in the expected forward direction, leading to poststimulus model updating due to the unanticipated event. To test this assumption, we used a multiclass linear discriminant analyser (LDA) which we trained on forward transition trials of the ordered condition in order to capture any prediction related neural activity. Afterwards, the classifier was tested on self-repetition trials to decode sound frequencies from brain activity in a time-resolved manner, providing classifier decision values for every sound frequency which were then transformed into corresponding transitions (e.g. 1_(t)_|1_(t-1)_ → repetition, 1_(t)_|4_(t-1)_ → forward,…). This was done for repetition trials of the ordered and the random entropy condition separately (see **Fig. 1B**). Resulting classifier decision tendencies (i.e. transformed decision values) for a forward transition versus a repetition were contrasted for every time-point from -0.3 s to 0.3 s of repetition trials (see **Fig. 1C**). Then, the average forward-vs.-repetition tendency was zero-centered and compared between entropy conditions (tested on ordered vs. tested on random repetitions). Since the classifier is trained on forward transitions only, bottom-up representations of the preceding sound are also labeled accordingly (carry-over classification effect). By balancing the transitions of both conditions, we ensured an equal carry-over effect across conditions which enabled us to isolate prestimulus prediction effects by contrasting “ordered” vs. “random”. Thus, we quantified “prediction tendency” as the classifiers prestimulus tendency to a forward transition in an ordered context exceeding the same tendency in a random context (see **Fig. 1C**). Analogously, we further quantified “updating tendency” as the poststimulus decision tendency towards a repetition transition (again for ordered > random). Using the summed difference across prestimulus/ poststimulus time, one prediction/ model updating value can be extracted per individual subject. Note that condition contrasting was also applied to ensure that individual differences in prediction tendency cannot be explained by subject specific signal-to-noise ratio (as no vacuous differences in SNR is to be expected between conditions). In addition, we used frequency decoding accuracy as a control variable to correlate with speech tracking variables of interest (see Appendix).

**Fig.1:**
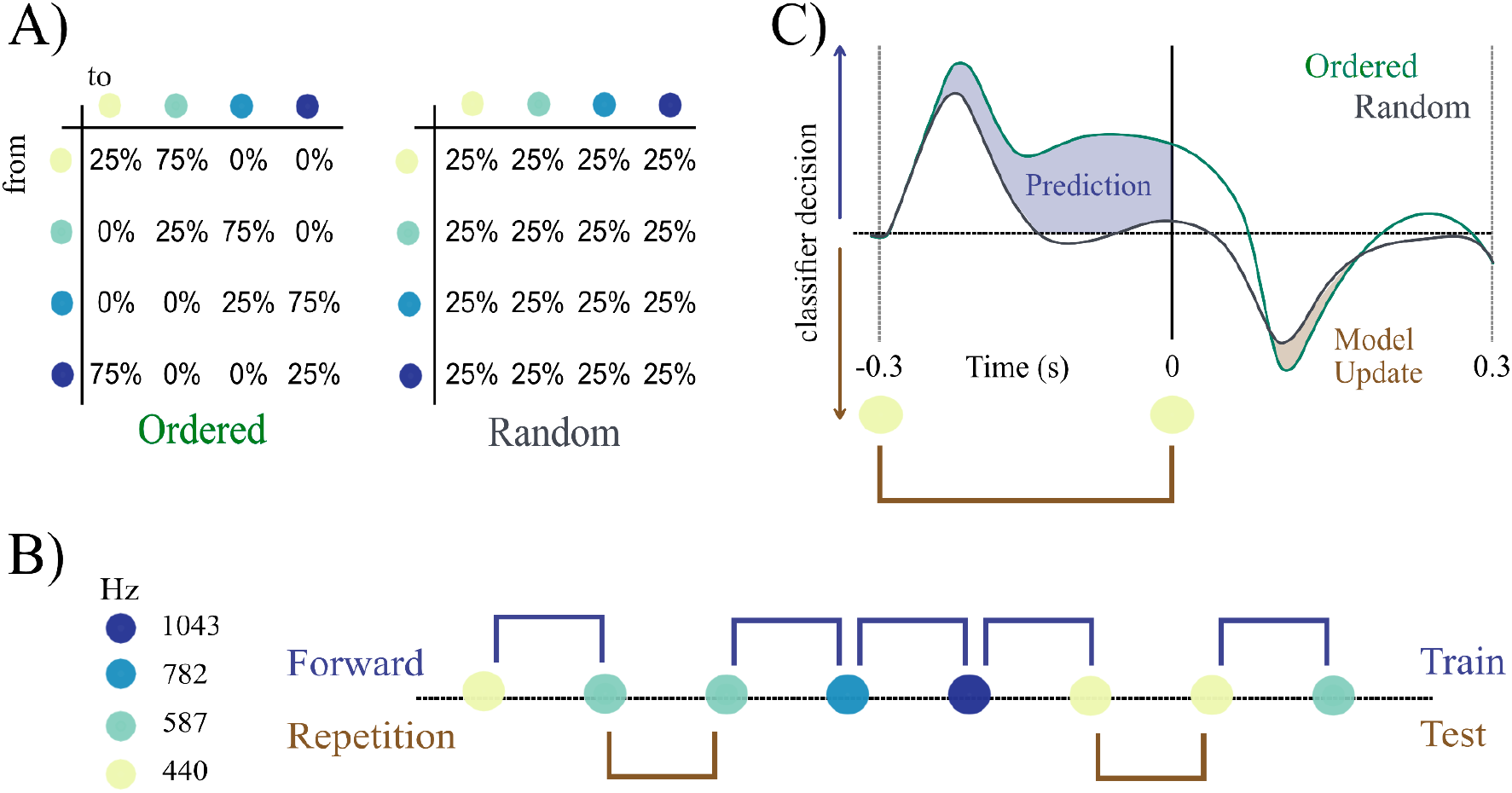
Quantification of individual prediction tendency. **A)** Participants have been presented with sequences of 4 different pure tones at a rate of 3 Hz. Transitional probabilities varied according to two different entropy conditions (ordered vs. random). **B)** Example of an ordered sound sequence. An LDA classifier was used to decode sound frequency from brain activity across time, trained on ordered forward transition trials and tested on all repetition trials. **C)** Expected classifier decision values contrasting the brains prestimulus tendency to predict a forward transition and poststimulus correction for the actually presented repetition. Individual prediction and model updating quantification result from the difference between conditions (ordered > random).

### Encoding of acoustic features

First, the speech envelope was extracted from the audio files of the target speaker using the Chimera toolbox (Smith et al., 2002) over a broadband frequency range of 100 Hz - 10 kHz (in 9 steps, equidistant on the tonotopic map of auditory cortex, see **Fig. 2B**). Afterwards, to quantify the neural representations corresponding to the acoustic envelope, we calculated a multivariate temporal response function (TRF) using the Eelbrain toolkit (Brodbeck et al., 2021). A deconvolution algorithm (boosting; (David et al., 2007)) was applied to the clear speech condition (0-dist) to estimate the optimal TRF to predict the brain response from the speech envelope for each individual virtual channel in source space (2982 voxels). The defined time-lags to train the model were from -100 ms to 500 ms. To evaluate the model, the 0-dist data was split into 4 folds, and a cross-validation approach was used to avoid overfitting (Ying, 2019). Furthermore, the same model was used to deconvolve speech envelope and brain data of the other two conditions (1-dist and 2-dist). The resulting predicted channel responses for all conditions were then correlated with the true channel responses in order to quantify the model fit and the degree of speech envelope tracking in a particular brain region.

**Fig. 2.:**
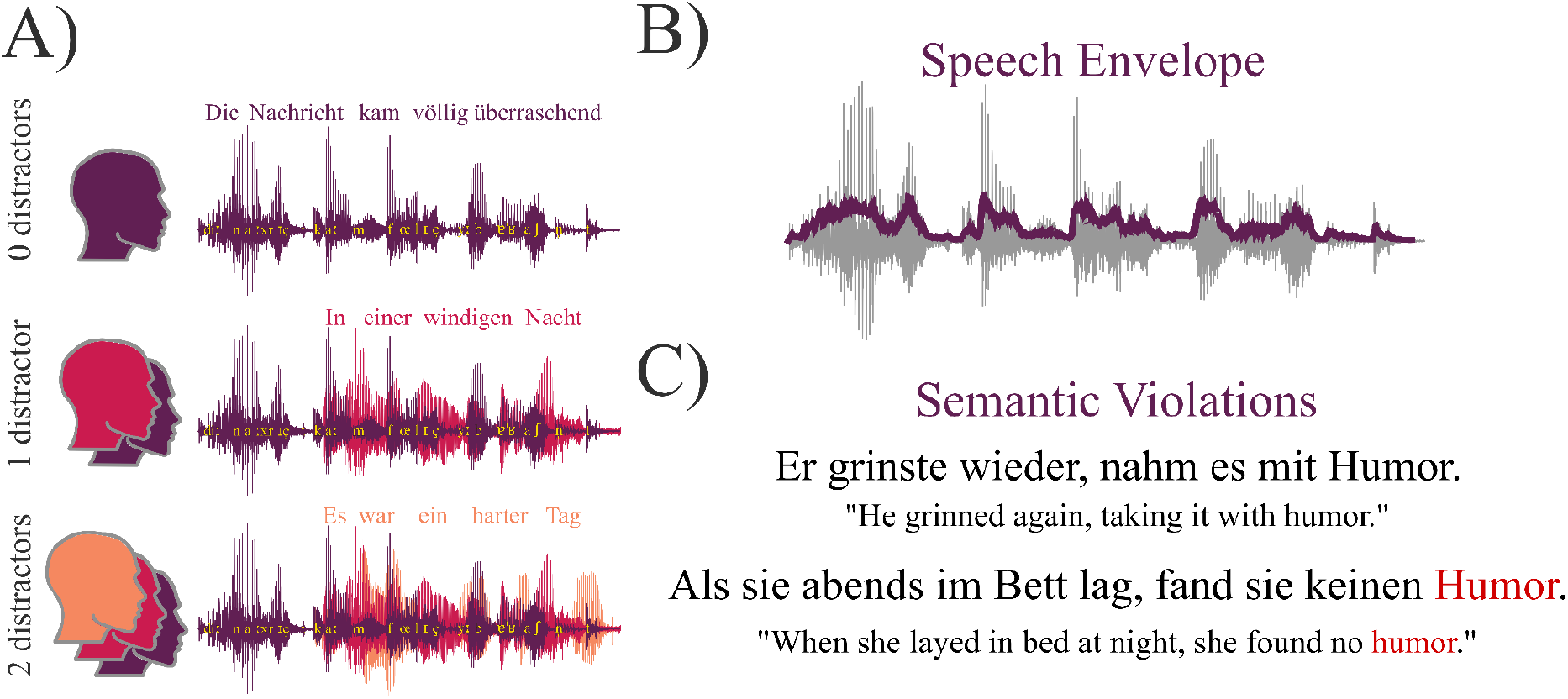
Multi speaker paradigm. A) Subjects were instructed to listen to a “target speaker” (purple). Depending on the experimental condition, the target speech stream was disrupted by one or two additional distracting speakers. B) The speech envelope of the target speaker was extracted and used to calculate a TRF (Brodbeck et al. 2021) with the associated brain activity recorded using MEG. C) Semantic violations were introduced randomly by replacing the last noun of a sentence with an improbable candidate to measure the effect of envelope encoding in conjunction with prediction tendency on the processing of semantic violations.

### Statistical analysis

To investigate the influence of individual prediction tendencies on speech tracking under different noise conditions, we used Bayesian multilevel regression models with Bambi (Capretto et al., 2022), a python package built on top of the PyMC3 package (Salvatier et al., 2016) for probabilistic programming. The correlation between predicted brain activity from speech envelope encoding and true brain activity was used as dependent variable, and separate models were calculated per voxel using the following formula according to the Wilkinson notation (Wilkinson & Rogers, 1973):

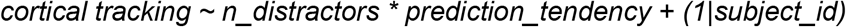

To investigate the influence of higher-level probabilistic structure of speech, we also calculated a model for which the dependent variable only included cortical tracking (i. e. speech envelope encoding) results of lexically identical nouns of high vs. low surprisal:

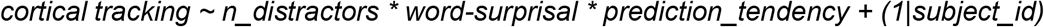

Before entering the models, prediction_tendency was zero-centered (note that in these interaction models other predictors are assumed to be zero to estimate effects of one predictor) and the number of distractors (0-2) was treated as a continuous variable. As priors we used the weakly-or non-informative default priors of Bambi (Capretto et al., 2022). For a summary of model parameters we report regression coefficients and 94% high density intervals (HDI) of the posterior distribution. From the HDIs we can conclude that there is a 94% probability that the respective parameter falls within this interval given the evidence provided by the data, prior and model assumptions. Effects were considered significantly different from zero if the HDI did not include zero. Further, it was ensured that for all models there were no divergent transitions (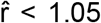 for all relevant parameters) and an effective sample size > 400 (an exhaustive summary of bayesian model diagnostics can be found in Vehtari et al., 2021). To further reduce the possibility of spatially spurious results, voxels were only considered as significant if a minimum of two significant neighbours could be detected. Subsequently, posterior distributions were aggregated over voxels that formed a cluster to report summary statistics for each effect.

Due to an error in the experimental code that resulted in loss of data on accuracy for true vs. false statements, we were not able to investigate listening comprehension on a behavioral level. To investigate the influence of background noise and individual prediction tendency on subjective perception we calculated two bayesian multilevel models using the following formulas:

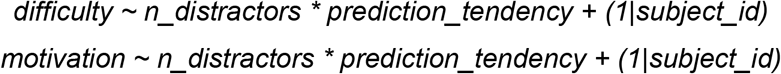

As dependent variables we used perceived task difficulty as well as self reported motivation (mean subjective ratings over blocks on a 5-point likert scale).

## RESULTS

### Quantifying individual prediction tendencies

First, we used an entropy modulation paradigm to quantify the individual prediction tendency which we define as the tendency to preactivate sound frequencies of high probability (i.e. a forward transition from one tone to another). Results for quantification of individual prediction tendencies are shown in **Fig. 3**. We find a clear prestimulus tendency to predict a forward transition during repetition trials in the ordered condition which exceeds the carry-over effect (see **Fig.3**). However, the data shows no visible difference in the correction for the actually presented repetition between an ordered and a random context. Model updating quantification was therefore dropped and only prediction tendency was used for further analysis.

**Fig.3:**
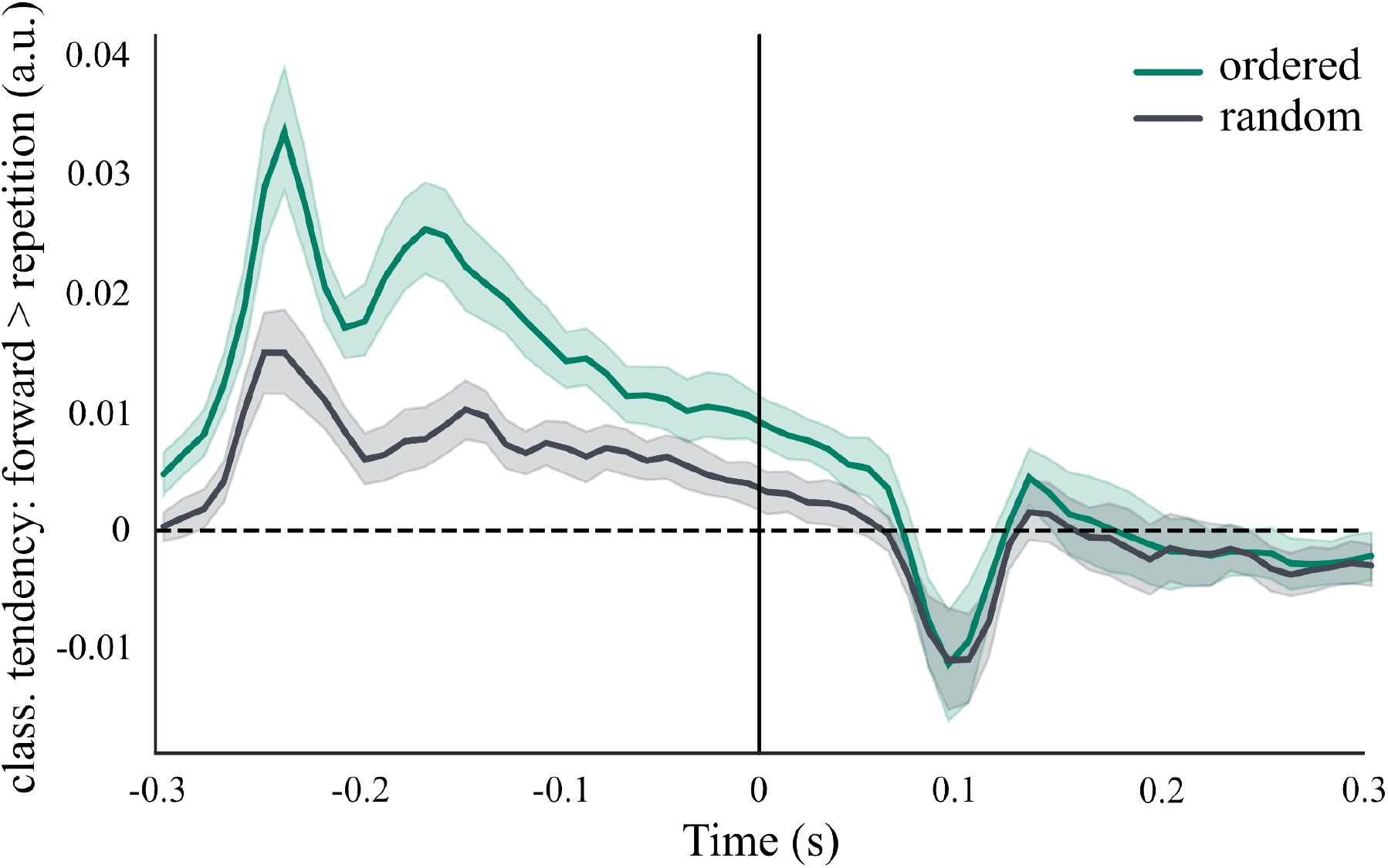
Individual prediction tendency. Time-resolved contrasted classifier decision: forward > repetition for ordered and random repetition trials. Classifier tendencies showing frequency-specific prediction for tones with the highest probability (forward transitions) can be found even before stimulus onset but only in an ordered context (shaded areas always indicate 95% confidence intervals).

### Speech comprehension becomes increasingly difficult with background noise

In order to investigate speech tracking, we used a multi speaker listening task. We find an effect for the number of distractors on self reported difficulty, indicating that the task is perceived more difficult as the number of distracting speakers increases. This result confirms that our modulation of the background noise achieves the desired effect. A more detailed description of all subjective task ratings (along with figures and summary statistics) can be found in the Appendix.

### Speech envelope encoding is modulated by individual prediction tendency and background noise

Across 6 continuous audiobook stories, we measured the extent to which the auditory speech envelope of a target speaker is encoded in 2982 virtual channels in source. First, we find a negative effect for the number of distractors on envelope encoding in bilateral auditory areas (b = -0.009, 94%HDI = [-0.023, -0.002], see **Fig. 4A**). This indicates that envelope encoding decreases with increasing noise for average prediction tendency. Second, we find a positive effect for prediction tendency in right perisylvian areas (namely inferior frontal, precentral and superior temporal cortex) and left inferior frontal cortex (b = 0.011, 94%HDI = [0.002, 0.025], see **Fig. 4B**). This suggests that individuals with stronger prediction tendency show an increased envelope encoding in these areas. There was also a negative effect for prediction tendency in the left inferior temporal cortex (b = -0.008, 94%HDI = [-0.015, -0.001]). Furthermore, our results indicate a positive interaction effect between the number of distracting speakers and prediction tendency in left inferior temporal and inferior frontal cortex (b = 0.004, 94%HDI = [0.001, 0.008]). This suggests that as speech gets more difficult to understand the role of prediction tendencies for envelope encoding increases in these left hemispheric areas (see **Fig. 4C**). However, we also find a negative interaction effect between the number of distracting speakers and prediction tendency in right inferior frontal, mid temporal and superior temporal areas (b = -0.005, 94%HDI = [-0.011, -0.001]). This indicates a right lateralized decreased effect for prediction tendencies on envelope encoding with increasing noise (see **Fig. 4C**). In sum, these results suggest an important, albeit differential role of individual prediction tendencies on speech envelope encoding.

**Fig. 4:**
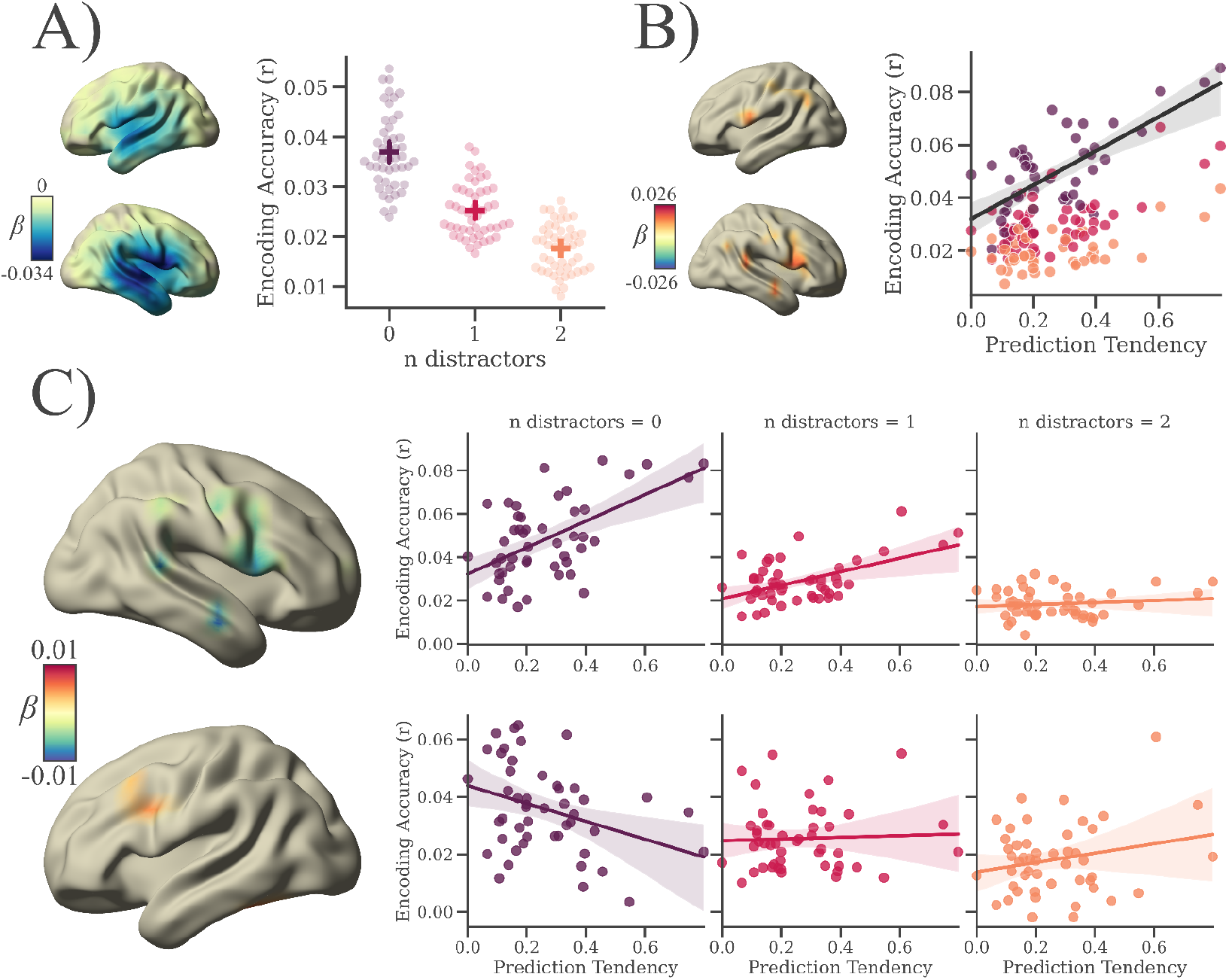
Speech envelope encoding is modulated by individual prediction tendency and background noise. A) The number of distracting speakers negatively affects speech envelope encoding, as background noise increases the encoding accuracy of the speech envelope decreases. B) Individual prediction tendency is positively associated with encoding of clear, continuous speech. C) However, as background noise increases, we notice a lateralized effect of prediction tendencies on speech envelope encoding. In right temporo-frontal areas the role of prediction tendencies on speech envelope encoding decreases with noise, while the opposite pattern was visible in left inferior frontal areas. In sum, these results suggest an important, yet differential role for the relationship between speech envelope encoding and individual prediction tendency.

### Violations of semantic probability differentially affect speech tracking

In order to investigate how the beneficial effect of a strong prediction tendency on speech processing in general might be affected by unpredictable events, we also induced rare semantic violations into our paradigm. In a direct comparison of lexically identical words, which have been differentially embedded into the semantic context, we find a positive effect on envelope encoding for words of high surprisal vs. low surprisal in left perisylvian cortex (b = 0.024, 94%HDI = [0.005, 0.047], see **Fig. 5A**). We also find a weak negative cluster showing a decreased encoding of surprising words in bilateral supplementary motor areas (b = -0.017, 94%HDI = [-0.031, -0.002], see **Fig. 5A**). For prediction tendency, again there was an overall positive effect in right superior temporal as well as bilateral inferior frontal cortex for encoding of words of low surprisal (b = 0.020, 94%HDI = [0.004, 0.041]). Furthermore, there was a left lateralized positive interaction between word surprisal and prediction tendency, indicating an increase in encoding of surprising words with increased prediction tendency in left angular gyrus (b = 0.017, 94%HDI = [0.004, 0.032], see **Fig. 5B**). Vice versa, we find a right lateralized negative interaction in mid occipital and superior frontal cortex, indicating that, with increased prediction tendency, encoding of surprising words is decreased whereas encoding of unsurprising words seems to be increased in these areas (b = -0.018, 94%HDI = [-0.033, -0.003], see **Fig. 5B**). Additionally, our results also show an effect for the number of distracting speakers along with several interaction effects with noise which are all listed in table 3 (see Appendix). In sum, the effects for the number of distractors for words of low surprisal replicate those of the previous section. We also find that effects of word surprisal along with the aforementioned interaction effects seem to diminish with increasing noise. We therefore conclude that the interaction between the influence of word surprisal and individual prediction tendency on encoding is negligible under conditions of high background noise. Instead, we focus our interpretation on the interaction effects of word surprisal and prediction tendency with respect to the clear speech condition.

**Fig. 5:**
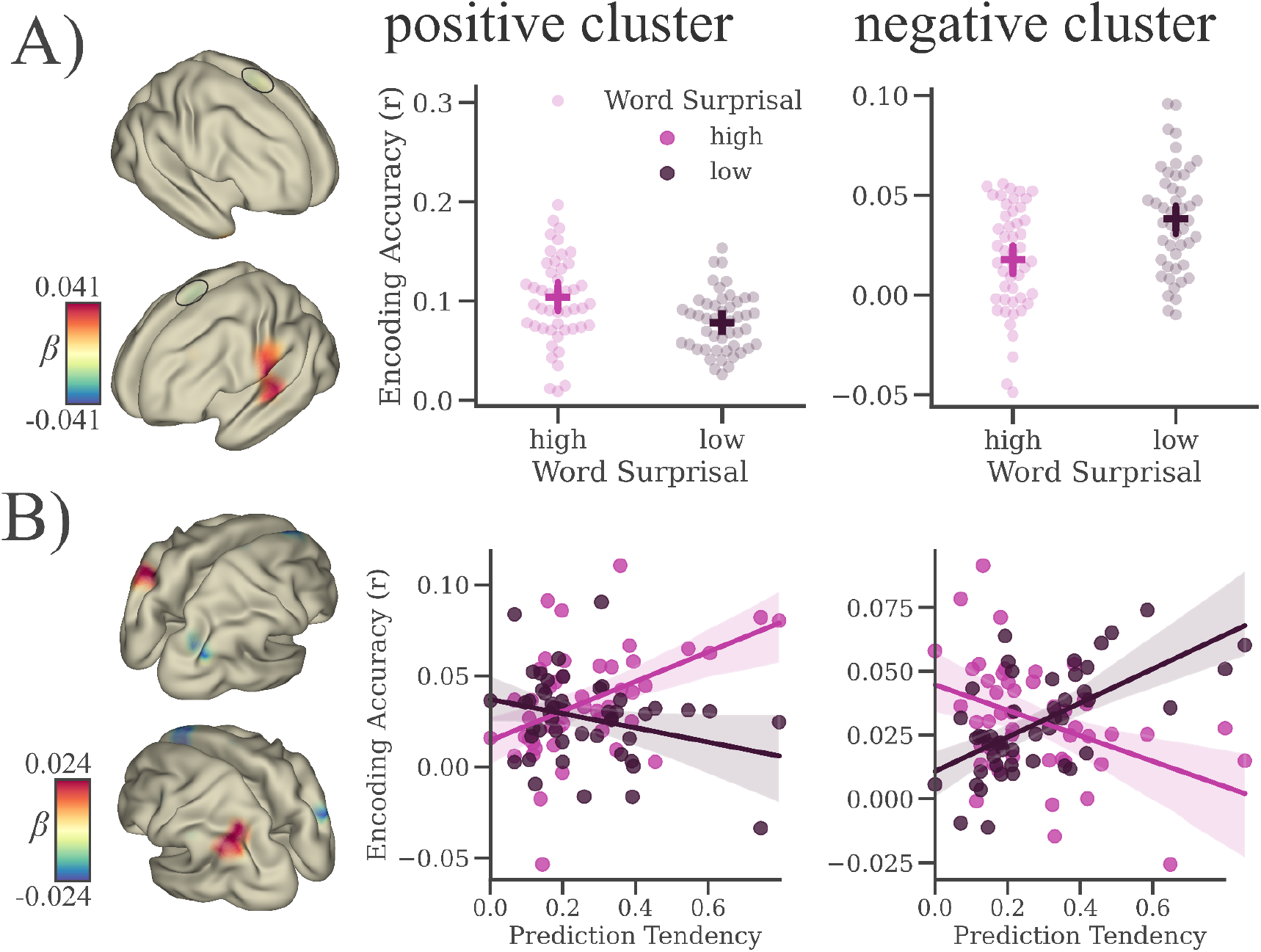
Violations of semantic probability differentially affect speech tracking. A) Lexically identical words differently affect the encoding of the envelope, if they are uttered in an (un-)related context. In the left temporal lobe we notice a positive association between word surprisal and encoding accuracy (the speech envelope was better encoded when words were uttered in an unrelated context). However, there was also a weak negative effect of word surprisal on envelope encoding in right bilateral motor areas (the speech envelope was better encoded when words were uttered in a related context; marked by a black circle). B) Furthermore, there was an interaction effect between individual prediction tendency and word surprisal, suggesting a decreased encoding of semantic violations in the right, versus an increased encoding of semantic violations in the left hemisphere, that scales with prediction tendency.

## DISCUSSION

Based on previous research emphasizing the ubiquity of the predictive brain, the aim of the current study was to investigate the supportive role of strong prediction tendencies in everyday situations such as listening. We propose a link between individual auditory prediction tendency and differences in the processing of spoken language. Across two independent paradigms, we connected the individual tendency to preactivate expected sound frequencies to the individual encoding of continuous narrative speech under varying conditions of background noise. Our results demonstrate that individual prediction tendency generally facilitates speech tracking. Furthermore, manipulating contextual predictability of speech by including rare semantic violations allowed us to investigate how this in itself beneficial effect is affected by unpredictable events, showing spatially differentiable interactions.

### Speech tracking is facilitated by individual prediction tendencies

Prediction tendency is conceptualized as an individual trait, a tendency that varies considerably across individuals and generalizes across situations, and that could be linked to psychiatric disorders such as autistic spectrum disorder (Pellicano & Burr, 2012) and schizophrenia (Corlett et al., 2019). Taking into account its explanatory value for such long-term and highly individualized disorders, predictions have worked their way from feedback connections carrying expected lower-level neural activity (Rao & Ballard, 1999) to an individual disposition that forms one of many core dimensions of personality. Here we make no such claim, but we strongly suggest that more emphasis is put on the investigation on the intraindividual stability of prediction tendencies (see Siegelman & Frost, 2015 for similar efforts regarding statistical learning). However, so far the beneficial aspects of prediction tendencies in complex listening situations have received little scientific attention.

Our findings show, for the first time, that the tendency to anticipate low-level acoustic features according to their contextual probability facilitates the cortical tracking of the speech envelope. Tracking of information conveyed by the speech envelope is critical for speech comprehension and the processing of low-level linguistic features (Schmidt et al., 2021; Vanthornhout et al., 2018). This suggests a potentially supportive role of the tendency to anticipate low-level acoustic features on the processing of naturally spoken language. Expectedly, this effect is most prominent in areas associated with auditory processing and speech perception such as inferior frontal as well as temporal regions of the right hemisphere, but also in Broca’s area of the left cortex (see **Fig. 4B**). To our knowledge, this is the first study to link speech tracking capabilities to individual differences in prediction tendency. This finding becomes all the more remarkable as our quantification of prediction tendency stems from a very different auditory paradigm that was completely independent of speech processing. Therefore, we can infer some generalizability of individual prediction tendencies across different listening situations. Most importantly, we argue that this link cannot be explained by individual signal-to-noise ratio or other attributes (such as linearity and distributive properties) that are known to have an influence on the performance of classification and encoding algorithms. First, prediction tendency is quantified as the difference in anticipatory predictions between conditions (thus uninformative idiosyncrasies in feature decodability are canceled out) and second, we find that individual decoding accuracy itself cannot predict envelope encoding, and its assumed beta shows no overlap with that of prediction tendency (see Appendix).

### The influence of prediction tendency on speech tracking decreases in the right and increases in the left hemisphere as background noise increases

Regarding the question on how the beneficial effect of prediction tendency on speech envelope tracking is affected by background noise, we find evidence for a dualistic effect showing a clear lateralization. More precisely, we find that the extent to which prediction tendency can explain speech tracking within the right hemisphere decreases with an increasing number of distracting speakers. Thus showing the strongest effect of prediction tendency on speech tracking whilst speech is clearly understandable. When simultaneously confronted with multiple speakers at a real cocktail party, the distracting speech streams are often spatially separable from the target stream, but in situations when a spatial segregation is not possible (for example in an online meeting) speech intelligibility is heavily decreased. As in our experiment the target and the distractor streams were both presented at phantom center, we created a similar situation. It is possible that with decreased intelligibility in bottom-up input, predictive processes are impaired from distortions in the auditory feedback-loop. The current results support our assumption that differences in prediction tendency can explain the variability in listening performance under optimal conditions.

However, we also find a left frontal cluster that shows decreased encoding of clear speech for individuals with stronger prediction tendencies. Furthermore, the negative association disappears and even slightly reverses with increasing noise. Thus, the extent to which the left inferior frontal cortex is engaged in speech envelope tracking in individuals with lower vs. stronger prediction tendencies seems to be dependent on background noise. For individuals with low prediction tendency we find substantial encoding of clear speech, whereas for individuals with strong prediction tendency speech encoding emerges only in conditions of increased background noise. One possible explanation is that differences in prediction tendency may result in differential allocation of attention depending on individual listening effort. For example, Lesenfants and Francart (2020) found that attention plays a crucial role in the cortical encoding of speech and showed an advanced encoding in frontal & fronto-central electrodes with focal attention. So far, differences in challenging listening situations with low signal-to-noise ratio (such as in a cocktail party situation) have often been linked to individual differences in selective attention (Kerlin et al., 2010). Yet, we argue that a predictive theory of individual listening experience is not exclusive but rather complementary to that as predictions are assumed to generate a selective attentional gain to the expected sensory feature (e.g. Marzecová et al., 2018). It seems possible that people allocate attention in a different or more parsimonious way depending on their tendency to rely on predictions. An alternative (though not mutually exclusive) interpretation is that individuals with weaker auditory prediction tendency have to rely more on sensorimotor integration in speech processing.

The general idea behind the concept of sensorimotor integration is that auditory processing is critically involved in speech production and, vice versa, that motor processes have a modulatory influence on speech perception. An integrative model was proposed by Hickok and colleagues (2011) in which the motor system promotes auditory forward predictions as part of a sensory-to-motor feedback circuit. Critically, they claim that the motor-speech networks are also involved in passive listening as feedback from auditory speech information, both self and others’, is relevant for production. Furthermore, it is assumed that forward predictions from the motor systems can also have perceptual consequences as they generate a sensory prediction that results in selective attentional gain for expected features. Based on this, we propose the following interpretation of our results: Individuals with a stronger tendency to generate purely auditory predictions are less reliant on predictions from the motor system during clear speech perception. However, as background noise increases, they start to use this mechanism as an alternative strategy to compensate for poor auditory input. The idea that the tendency for multisensory integration shows great interindividual variability is not new but has so far mainly focused on audiovisual integration for speech perception (e.g. Nath & Beauchamp, 2012). We suggest that future research should explore the possibility of an interaction between individual prediction tendency and sensorimotor integration during speech perception in challenging listening situations.

### Violations of semantic probabilities interact with individual prediction tendencies

To test the influence of individual prediction tendencies on changes in semantic probabilities during speech processing, we compared speech envelope encoding between a subset of surprising target words that are semantically unrelated to the preceding context and their lexically identical counterparts for which contextual predictability was not manipulated. Our results show an increased envelope tracking for surprising words in the left perisylvian cortex. Over the last decades, most research that has investigated the effects of surprise in speech perception has focused on event-related potentials and the typical N400 component - a negativity at 300 – 600 ms in central and/or parietal regions. It is a well established finding that unexpected stimuli raise a stronger response (or from another perspective: expected stimuli evoke a reduced neural response), but the relative information content of these responses is still debated (de Lange et al., 2018). Our findings show considerable spatial overlap with previous studies that have focused on surprisal-evoked responses in speech perception (Frank & Willems, 2017; Willems et al., 2016), complementing them in two ways. First, we provide evidence that these responses are not driven by a rather unspecific error signal but instead actually encode information about the surprising stimulus itself. Our results indicate that the low-level acoustic representation is sharpened for high compared to low surprisal and, importantly, that this sharpening is a consequence of contextual unpredictability and cannot be explained by any differences in bottom-up input. Second, even though the impact of predictions on speech perception has widely been emphasized (e.g. Heilbron et al., 2021), the question of how individual prediction tendency interacts with the processing of semantic violations has, to the best of our knowledge, not been investigated so far. Originally, we expected a disrupted encoding of surprising words, scaling with individual prediction tendency. Instead, we find a spatially dissociable interaction effect, indicating the expected interference for surprising words in right occipital and superior frontal regions on the one hand (or hemisphere), but also an increased encoding of surprising words in left angular gyrus with increased prediction tendency on the other. These findings allow for different interpretations and in the following the two interaction effects are separately discussed in more detail.

The superior frontal and occipital locations, for which we find a negative impact of surprise and prediction tendency, could suggest a possible contribution of eye movements to the encoding of acoustic information. Eye movements have already been linked to top-down auditory attention, even in the absence of any visual stimulation, and great interindividual variability has been observed in the extent to which they are used to support auditory attention (Braga et al., 2016). As the current study focused on qualitative differences (i.e. the extent to which an auditory feature is represented) rather than quantitative differences in power or amplitude, we currently do not rule out the general possibility that speech envelope may also to some extent be tracked via eye-movements.

In contrast, we also find a positive interaction of prediction tendency and word surprisal in the left angular gyrus. One possible interpretation is that this enhanced encoding reflects a prediction error that is increased in individuals with stronger prediction tendency. Prediction errors are generally assumed to convey unexpected information to update internal models and improve future predictions, however their content and functionality can vary across situations and brain regions (Den Ouden et al., 2012). From the current results we cannot draw any conclusions on internal model updating, but we suggest further investigations on the impact of error-driven learning in speech perception with respect to interindividual differences.

## Limitations

There are several limitations to the interpretation of our results. First, we were not able to directly link prediction tendencies to differences in behavioral task performance (due to data loss) or to subjective listening experience. Although one might argue that these variables are of considerable importance for real-life implications, in the current study they are likely to be biased by floor/ceiling effects (see Appendix). Second, we did only focus on one acoustic feature although speech perception encompasses many more linguistic features (such as formants, phonemes, syllable rate etc.), and even though we demonstrated an effect of word surprisal on low-level encoding, we did not investigate the tracking of such higher-level features directly. Future research should focus on the explanatory value of individual prediction tendency for speech processing across all levels along the linguistic hierarchy.

## Conclusion

We support the assumption that predictive processing is an ubiquitous perceptual phenomenon and that it is crucial for continuous speech perception. Furthermore, we argue that the tendency to engage in these predictive processes is different from individual to individual and that this variability can explain differences in listening experience. Importantly, individual prediction tendency can be assessed independently of the speech perception it facilitates, thus we can infer some generalizability of auditory predictions across different listening situations.

## ACKNOWLEDGEMENTS

J.S. and Q.G. are supported by the Austrian Science Fund (FWF; Doctoral College “Imaging the Mind”; W 1233-B). Q.G. is also supported by the Austrian Research Promotion Agency (FFG; BRIDGE 1 project “SmartCIs”; 871232) and F.S. is supported by WS Audiology. Thanks to the whole research team. Special thanks to Tom Westfeld for helping us with the audio recordings and to Manfred Seifter for conducting the MEG measurements.

## APPENDIX

### Stimulus Material

We used audio recordings of excerpts from German books and short-stories (listed in the table below) for our multispeaker listening task. Excerpts were selected prior to recording in such a way that one trial had an approximate reading duration of 3 - 4 min (mean = 3.46 min, range = 3.05 - 4.07 min). In total 27 trials were recorded from the following material:

#### Speech tracking effects cannot be explained by differences in pure tone decoding

To show that the relationship between prediction tendency and the encoding of speech features cannot merely be explained through individual differences in signal-to-noise ratio we added individual (zero-centered) pure-tone decoding accuracy peaks from the entropy modulation paradigm into our model. Envelope encoding was averaged over voxels that showed a significant effect of prediction tendency and used as dependent variable. Model summary statistics (see Table 2) show that individual pure-tone decoding cannot predict envelope encoding results (b = -0.001, 94%HDI = [-0.004, 0.002]) and the posterior probability distribution shows no overlap with the posterior probability distribution of prediction tendency (b = 0.011, 94%HDI = [0.007, 0.014]). We therefore conclude that the influence of individual prediction tendency on speech encoding cannot be attributed to differences in decoding results per se.

**Table 1:** List of all ebooks and short stories that were used as basis for audio material

|  |
| --- |
| “Ein zauberhafter Schrebergarten” by Anja Pompowsk<sup>1</sup> |
| “Winterzauber in der kleinen Keksbäckerei” by Holly Hepburn<sup>1</sup> |
| “Darius’ Radius” by Maida Thesy<sup>1</sup> |
| “Die Möwe Jonathan” by Richard Bach<sup>1</sup> |
| “Die Eskimos: Geschichte und Schicksal der Jäger im hohen Norden by T. Jeier<sup>1</sup> |
| “Sofies Welt” by Jostein Gaarder<sup>1</sup> |
| “Die zwei Weiden” and “Der Zaunkönig und die Rose” by Florian C. Pichler<sup>2</sup> |
| “Der Goldstrauch und 66 weitere Kurzgeschichten für zwischendurch” by Alfred Bekker<sup>2</sup> |
| “Der Schneesturm” by Nadja Rohner<sup>2</sup> |
| “Die Burg” by Werner Kistler<sup>2</sup> |
| “Kishon's schönste Geschichten für Kinder” by Ephraim Kishon<sup>2</sup> |

Note: <sup>1</sup>main stories narrated by a target speaker (3 x ~3min excerpts each), <sup>2</sup>distractor stories narrated by a distractor speaker (18 x ~3min excerpts in total)

**Table 2:**
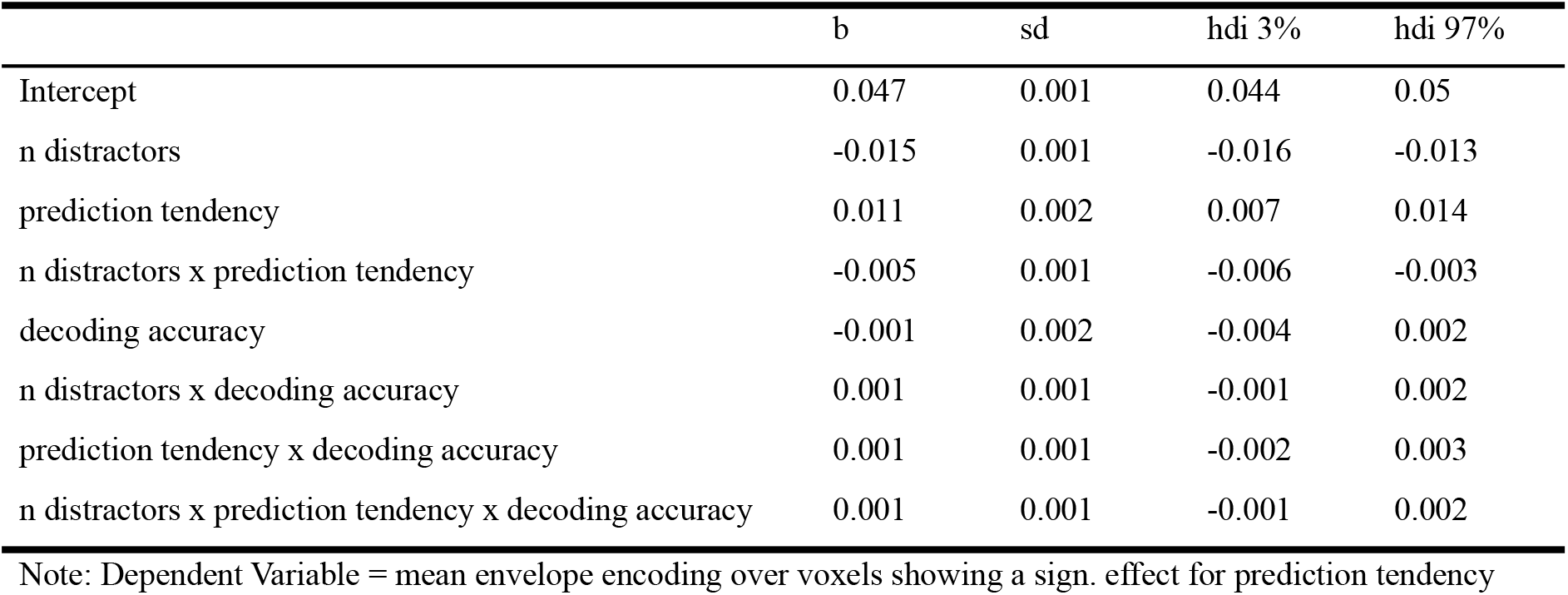
Model summary statistics including pure-tone decoding accuracy

|  | b | sd | hdi 3% | hdi 97% |
| --- | --- | --- | --- | --- |
| Intercept | 0.047 | 0.001 | 0.044 | 0.05 |
| n distractors | -0.015 | 0.001 | -0.016 | -0.013 |
| prediction tendency | 0.011 | 0.002 | 0.007 | 0.014 |
| n distractors x prediction tendency | -0.005 | 0.001 | -0.006 | -0.003 |
| decoding accuracy | -0.001 | 0.002 | -0.004 | 0.002 |
| n distractors x decoding accuracy | 0.001 | 0.001 | -0.001 | 0.002 |
| prediction tendency x decoding accuracy | 0.001 | 0.001 | -0.002 | 0.003 |
| n distractors x prediction tendency x decoding accuracy | 0.001 | 0.001 | -0.001 | 0.002 |

Note: Dependent Variable = mean envelope encoding over voxels showing a sign. effect for prediction tendency

**Table 3:** Model summary statistics for comparison of high vs. low word surprisal

|  |  | b | sd | hdi 3% | hdi 97% |
| --- | --- | --- | --- | --- | --- |
| Intercept | L+R: auditory C. | 0.046 | 0.024 | 0.013 | 0.101 |
| n distractors | L+R: auditory C. | -0.019 | 0.010 | -0.043 | -0.005 |
| prediction tendency | L+R: front.inf./ precentr., R: temp.sup. | 0.020 | 0.010 | 0.004 | 0.041 |
| word surprisal (high > low) | L: postcentr., temp.sup., perisylv.A. | 0.024 | 0.011 | 0.005 | 0.047 |
|  | L+R: SMA | -0.017 | 0.008 | -0.031 | -0.002 |
| n distractors x word surprisal | L: SMA, S.calc., R: temp.mid., occ.mid. | 0.015 | 0.006 | 0.003 | 0.027 |
|  | L: postcentr., R: front.mid. | -0.016 | 0.007 | -0.031 | -0.003 |
| word surprisal x pred. tend. | L: pariet.inf., angular G. | 0.017 | 0.007 | 0.004 | 0.032 |
|  | R: front.sup.med., occ.mid. | -0.018 | 0.008 | -0.033 | -0.003 |
| n distractors x pred. tend. | L: occ.sup. | 0.010 | 0.005 | 0.002 | 0.019 |
|  | L: precentr., R: temp.mid./sup., front.mid. | -0.012 | 0.005 | -0.023 | -0.002 |
| n distractors x word surprisal x prediction tendency | L: front.sup.med., pre-/postcentr., R: occ.mid | 0.014 | 0.006 | 0.003 | 0.027 |
|  | L: perisylv.A., pariet.inf., R: front.inf./mid. | -0.015 | 0.007 | -0.028 | -0.003 |

Note: Dependent Variable = mean envelope encoding over voxels showing a sign. effect for each predictor

#### Violations of semantic probabilities interact with individual prediction tendencies as well as with background noise level

As we investigated how semantic violations are encoded differently in comparison to their lexical identical counterparts we further included individual prediction tendency as well as the number of distractors into our model. The results can be found in the table below. Note that for most predictors we find a positive as well as a negative effect albeit in spatially different locations (labels for all voxels were obtained using a template atlas; Tzourio-Mazoyer et al., 2002).

#### Behavioral Results

As we were not able to investigate speech comprehension on a behavioral level we had to use self reported ratings on difficulty (on a 5-point likert scale) as a proxy for individual perception. We clearly find an effect for the number of distractors indicating that the task was perceived more difficult with increasing background noise (b = 1.251, 94%HDI = [1.173, 1.325]). There was no visible effect for individual prediction tendency (see **Fig. 6A** and Table 4) indicating that across subjects the task was perceived equally difficult across subjects with different prediction tendencies. It should be noted, however, that the ratings for the dist-0 condition and the dist-2 condition show a pronounced floor and ceiling effect respectively (see **Fig. 6B)**. Further we investigated whether background noise and/or individual prediction tendency had an effect on motivation. We found no difference in self reported motivation between conditions (b = -0.059, 94%HDI = [-0.124, 0.005]) and no effect for individual prediction tendency on motivation (b = -0.096, 94%HDI = [-0.318, 0.116]; see **Fig. 6B** and Table 5).

**Table 4:**
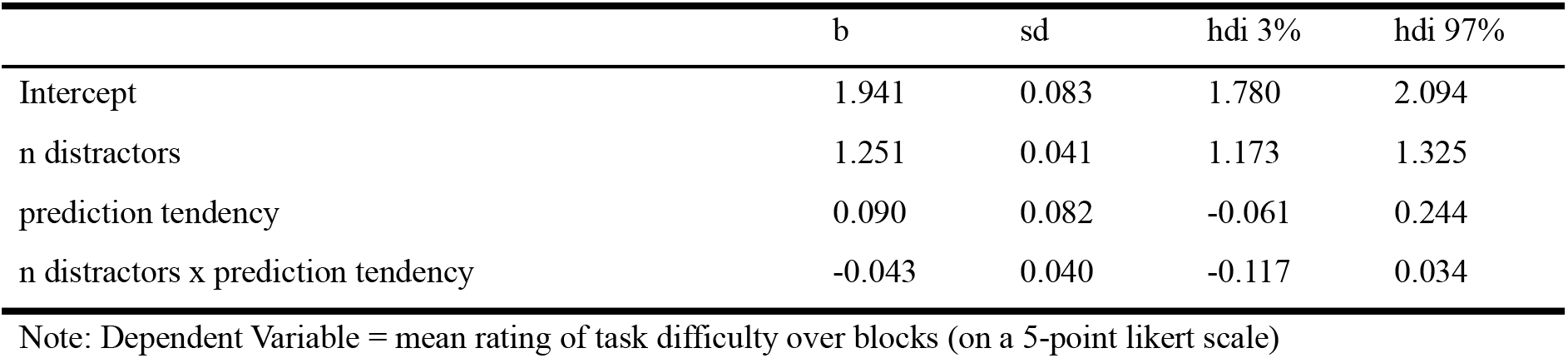
Model summary statistics for perceived task difficulty

**Table 5:** Model summary statistics for self reported motivation

|  | b | sd | hdi 3% | hdi 97% |
| --- | --- | --- | --- | --- |
| Intercept | 3.889 | 0.120 | 3.659 | 4.108 |
| n distractors | -0.059 | 0.035 | -0.124 | 0.005 |
| prediction tendency | -0.096 | 0.118 | -0.318 | 0.116 |
| n distractors x prediction tendency | -0.030 | 0.035 | -0.097 | 0.034 |

Note: Dependent Variable = mean rating of motivation over blocks (on a 5-point likert scale)

**Fig.6:**
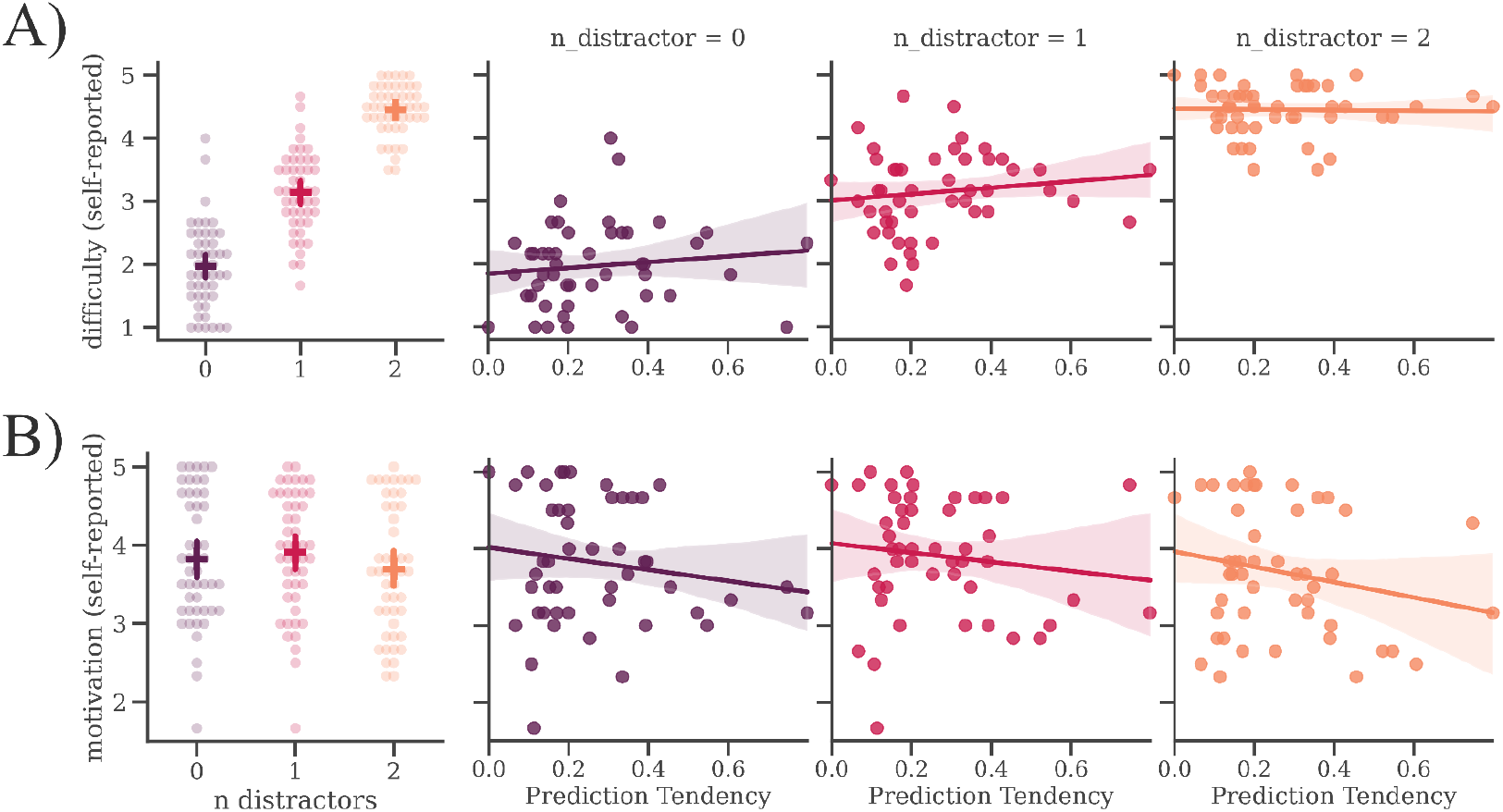
Background is affecting subjective ratings of difficulty in a multi-speaker listening task. A) Perceived task difficulty (indicated by subjective ratings on a 5-point likert scale) increases with the number of distractors and is not affected by individual prediction tendency. B) Motivation (indicated by subjective ratings on a 5-point likert scale) is neither affected by the number of distractors nor by individual prediction tendency.

